# Shape and rate of landscape change trajectories influence species persistence

**DOI:** 10.1101/2025.08.04.666294

**Authors:** Nivedita Varma Harisena, Adrienne Grêt-Regamey, Maarten J. van Strien

## Abstract

Species persistence in changing landscapes is shaped not only by present-day habitat conditions but also by the historical trajectories of habitat loss and fragmentation *per gradus*. Using a simulation-based framework, we investigated how long-term patterns of landscape change, defined by initial and final habitat area and rate of change of landscape configuration, affect species metapopulation persistence. Using 3,600 simulated landscape change scenarios with varying initial and final habitat conditions, we coupled dynamic habitat loss with individual-based metapopulation models to assess time till extinction under different rates of landscape change, scaled to species generation time. Our results show that while final habitat amount strongly influences persistence under rapid change, in slower-changing landscapes, species persistence is substantially modulated by historical habitat extent and the trajectory of landscape configuration. Effective mesh size (MESH) and its rate of change was a stronger predictor of persistence in scenarios with high initial habitat. In contrast, the number of patches (NP) was more influential in already degraded or low-change scenarios, where final configuration closely aligned with initial conditions. Notably, we observed long extinction delays in landscapes that began with high habitat area and connectivity but became fragmented gradually, highlighting that historical continuity can mask imminent extinction risk in current fragmented systems. Crucially, species persistence correlated most strongly with the *rate* of change in MESH and NP when scaled to species generation time, revealing that extinction lags are driven by how quickly habitat configuration deteriorates relative to species’ life cycles. These findings underscore the need to incorporate long-term changes of spatial configuration into biodiversity assessments, as present-day species presence may hold underlying extinction debts.

## 1. Introduction

In a changing world, species persistence represents a complex phenomenon in landscape ecology. In this context, persistence refers to the continued presence of species in metapopulations due to colonisations from nearby patches despite local extinctions (Hanski, 1998). In landscapes where this equilibrium between colonisation and extinction is not maintained (i.e. where colonisation rates do not allow for sufficient recovery in local patches), species are prone to extinction. Such extinction can sometimes be delayed; for example, colonisation events can continue to allow species persistence following disturbance, leading to extinction lagging behind habitat disturbance by a significant amount of time (Hanski and Ovaskainen, 2002). This delay accommodates species occurrences in landscapes, irrespective of the lack of capacity of the habitats to foster such occurrences. Such occurrences would continue to decrease in the future, regardless of the absence of additional disturbances to the landscape, and have been termed as an ‘extinction debt’ (Kuussaari et al., 2009). Accordingly, it is critical to understand which processes lead to such time-lagged extinctions and what the duration of the delay will be (Chen et al., 2023; Hylander and Ehrlén, 2013a). Understanding these processes can allow us to view current biodiversity within its spatio-temporal contexts, thereby aiding nature conservation strategies (Watts et al., 2020).

Species’ continued persistence in landscapes that have undergone habitat loss and fragmentation has been theorised in the seminal work on metapopulation dynamics by Hanski and Ovaskainen (2002). Most studies on time-lagged processes focus on short-term changes (Bertassello et al., 2021; Hanski and Ovaskainen, 2002; Johst et al., 2002) or only use snapshots of historical landscapes (Chen and Peng, 2017; Harisena et al., 2024; Herrault et al., 2016; Pan et al., 2022; Vellend et al., 2006). The latter studies aim to illustrate the presence of an extinction debt by identifying correlations between past habitat area and contemporary species diversity. However, recent studies increasingly recognise that extinction processes are not only governed by the historical and current states of habitats, but also by the shape of the complete trajectory of landscape change over time (Liao et al., 2022). Notably, few studies have examined the influence of long-term trajectories of landscape change on species persistence. Graham et al. (2017) identified the thresholds and patterns of habitat loss (e.g. fragmentation into small versus large patches or rare versus common patches) that shape species persistence, highlighting tipping points beyond which metapopulation rescue effects can no longer offset habitat loss. However, these authors did not consider the impact of continuously changing landscapes. Other studies have identified how stable versus declining habitats influence species persistence (Guardiola et al., 2013; Helm et al., 2006; Ridding et al., 2021). However, these studies did not differentiate between rates of landscape change. Such rates of change are hypothesised to serve as key determinants for delays in ecological responses (Lütolf et al., 2009; Williams et al., 2021). Thus, it is important to investigate the shapes of trajectories and rates of change to better understand the persistence of species in changing landscapes.

In addition to the shapes and rates of trajectories of landscape change, species traits such as dispersal ability and generation time may also influence the time lags over which metapopulations persist. For example, dispersal ability determines the spatial extent over which individuals can move between habitat patches, which can affect the viability of enabling potential ‘rescue effects’ over time in fragmented landscapes if connectivity is sufficient (Hanski and Ovaskainen, 2000; Loreau et al., 2003). Similarly, generation time critically influences the ability of species to respond to changes in the landscape. Species with short generation times may respond more quickly, while those with long generation times are more likely to accumulate extinction debts, especially when landscape change is gradual or prolonged (Watts et al., 2020). However, empirically identifying the influence of species traits on extinction time lags remains challenging due to the long temporal scales involved, confounding ecological processes and the scarcity of long-term datasets (Ridding et al., 2021). A study by Jiménez-Franco et al. (2022) addresses this by demonstrating that movement limitations in long-lived species can contribute to delayed extinction when tracking cumulative landscape changes from 1956 and projecting impacts up to 200 years into the future. Their findings suggest a propagation of time-lagged effects across demographic processes, although landscape change in their study was limited to three discrete historical time points within a specific case-study area. Other species traits that can also influence the time lag in extinction include the reproduction rate and intra-species competition—both of which can define the extinction rate of a species. The lower the extinction rate, the higher the likelihood of delayed extinction (Hylander and Ehrlén, 2013a). Thus, both landscape (habitat) parameters and species traits together define species persistence in changing landscapes.

Trajectories of landscape change that affect biodiversity can be defined in several ways, including changes in habitat quantity, habitat connectivity and the structure of the surrounding landscape matrix (Fletcher et al., 2024; Harisena et al., 2025). While the consequences of direct habitat loss are relatively well-established, the role of habitat fragmentation, particularly in terms of spatial configuration, has been the subject of ongoing debate (Miller-Rushing et al., 2019). Krauss et al. (2010) have shown that habitat fragmentation shows delayed impacts on species occurrence, while other studies have shown that such time-delayed effects depend on variables indicating the configuration of habitat patches (Herrault et al., 2016; Semper-Pascual et al., 2021b). Changes in patch configuration can disrupt dispersal routes, reduce effective habitat accessibility and alter population dynamics, particularly in species with limited mobility or specialised habitat requirements (Kuipers et al., 2021). Thus, in addition to the habitat area, changes in the configuration of habitat patches may also represent an important parameter in defining species persistence. To investigate this, landscape metrics that quantify both habitat quantity and configuration can be used (Jaeger, 2000; Justeau-Allaire et al., 2022).

This study aims to develop a theoretical understanding of the impact of long-term and continuous landscape change trajectories on species persistence. Specifically, we investigate how the shapes and rates of landscape change trajectories—not only sudden disturbances—influence the length of time that species persist in a landscape. We incorporate change in two ways: by directly modifying the landscape and by considering landscape change with reference to species generation times. This can provide a framework applicable to a wide range of species in a changing world. Using landscape change simulations, we address the question of species persistence in landscapes that have similar final habitat areas but different past landscape change trajectories in terms of habitat loss and configurational changes. Simulated landscapes serve as an appropriate alternative to empirical studies, enabling the identification of well-defined (controlled) change trajectories that incorporate a broad range of initial habitat areas and rates of change (Slager and De Vries, 2013; van Strien et al., 2016). We further simulate species metapopulation dynamics within these changing landscapes to estimate species persistence using individual-based models. To explore each trajectory of change, we further examine landscape metrics such as effective mesh size and the number of patches to understand their relative roles in species persistence. Through this approach, we describe patterns of landscape change trajectories associated with varied species persistence across different long-term landscape trajectories, offering a theoretical framework to guide ecological investigations of delayed species responses in dynamic landscapes.

## 2. Methods

We used a simulation-based approach to investigate how long-term trajectories of landscape change influence species persistence. Our framework couples dynamic simulations of landscape change with individual-based models of metapopulations to explore the interactions between landscape history and generation times in shaping time-lagged extinction responses.

### 2.1. Study design

The study followed four analysis steps. First, a series of initial landscapes was simulated using spatial correlation derived from Matérn variogram structures for a range of initial habitat areas (Fig. 1.a). The habitat area in these initial landscapes was then linearly reduced over 20 time steps until a specified final habitat area was reached. For various combinations of initial and final habitat area, a total of 3,600 landscape change series were simulated. For each trajectory, the change in effective mesh size (MESH) and number of patches (NP) was calculated to capture connectivity and fragmentation dynamics, respectively (Fig. 1b). Species metapopulation dynamics were then simulated on these landscapes using an individual-based metapopulation model with assumptions on species traits and dispersal (Fig. 1.c). Landscape change steps were integrated into the species simulation by updating the habitat conditions after several species generations, where each generation faced a sequence of reproduction, dispersal and mortality due to competition and lack of habitat. Species persistence, measured as the log-transformed mean global time to extinction, was derived from the individual-based models (Fig. 1.d; Tao et al., 2024). Making use of random forest models, we determined which properties of the landscape change trajectories influence species persistence. Predictor variables included the temporal trajectories of MESH and NP, assessed across all scenarios and stratified by specific final habitat percentages. Additionally, we applied time series clustering to investigate how different landscape trajectories influence species persistence. Finally, we disentangled the relative contributions of the rate of landscape change (effective slope) versus initial habitat conditions in explaining persistence outcomes.

**Figure 1:**
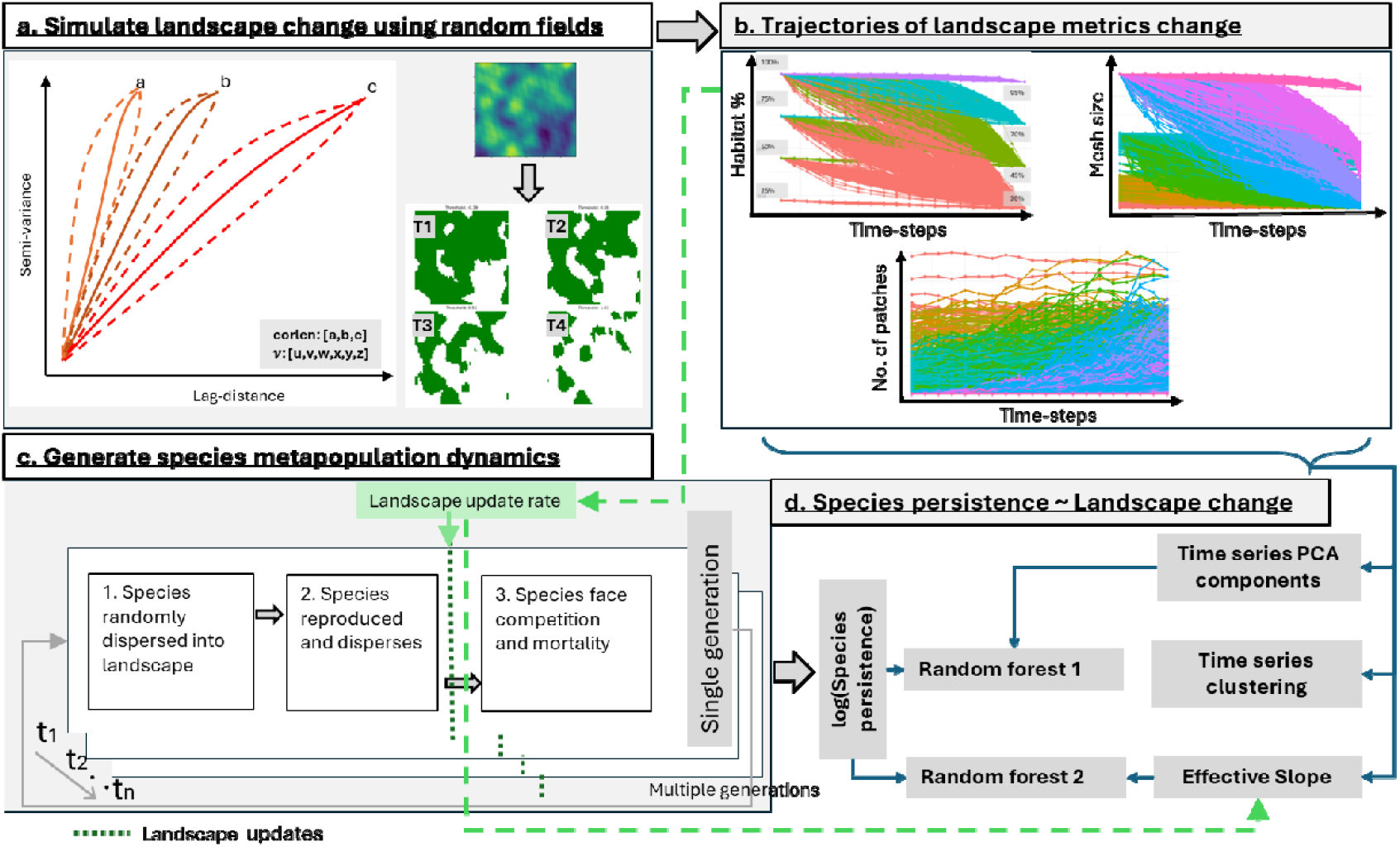
Overview of the workflow of the study. a) Simulating landscape change scenarios. b) Calculating the time series of landscape change. c) Generating species metapopulation dynamics. d) Relating species persistence to landscape metric trajectories, rates and initial conditions.

### 2.2. Landscape change simulations

To generate landscape change series, we generated Gaussian random fields of 60 by 60 pixels using Matérn covariance functions, varying the correlation length parameter (*corlen* 5, 10, 20) and a smoothness parameter (0.5, 1, 5, 10, 20) using the GStools package in Python (Müller et al., 2022). This produced a set of random landscapes with varying degrees of spatial autocorrelation, as illustrated in Fig. 1a and detailed in Supplementary Material S1. In spatial models using the Matérn variogram, the correlation length parameter determines the range over which values are spatially correlated. Higher correlation lengths result in larger, more contiguous random fields as spatial similarity persists over longer distances. Lower values produce more fragmented fields due to rapid spatial decorrelation. The smoothness parameter, denoted by ν (nu), controls the shape of the variogram near the origin: lower ν values lead to rougher fields with sharp local variation, while higher ν values generate smoother fields with more gradual spatial transitions. The mapped random field values were subsequently thresholded at multiple levels across their value distribution to produce binary habitat maps, where pixels exceeding a given threshold were classified as habitat and those below the threshold as the matrix. Threshold values were selected to linearly decrease the proportion of habitat across the field, creating a series of 20 habitat maps per random field. Since the underlying spatial structure of each random field remained fixed, the sequence of thresholded maps simulated a temporal trajectory of landscape change driven solely by habitat loss (see Supplementary Material S1 for details).

We assumed that all our simulated landscapes showed a certain level of habitat loss (minimally 5% loss and maximally 80% loss). The simulation design incorporated four initial habitat area levels (100, 75, 50 and 25%) and four corresponding final habitat area levels (95, 70, 45 and 20%), yielding 10 distinct habitat decline scenarios across different combinations of initial and final conditions (i.e. 100→95, 100→70, 100→45, 100→20, 75→70, 75→45, etc). For each scenario, 20 independent realisations were generated for each combination of *ν* and *corlen* parameter values, resulting in a total of 360 simulated time series per habitat decline scenario and 3600 time series across all scenarios.

### 2.3. Trajectories and metrics of landscape change

In this study, trajectories of landscape change were defined by initial and final conditions and rates of change in a) habitat area and b) the connectivity between the habitats defined by configurational changes. Habitat area and connectivity contribute differently to species time-lagged responses: larger historical areas allow for larger initial populations, which may have an effect on metapopulation dynamics under the process of habitat loss, whereas higher connectivity can facilitate the rescue of a locally extinct population from nearby patches that still have the capacity to host populations (Hanski, 1998). The initial and final area of the habitat patches was the controlled setting for the simulations, as shown in Fig. 1.b and Supplementary Material S2. To compute the configurational changes, we calculate different landscape metrics that additionally pertain to the changes in connectivity and fragmentation *per gradus* in the landscape. These configuration changes arise from the stochastic nature of the random field generation.

For each of the 3600 simulated landscape series, we calculated a temporal series of 12 landscape metrics quantifying connectivity and fragmentation using the *flsgen* package in R v4.4.1 (Justeau-Allaire et al., 2022) (see Fig. 1.b and Supplementary Material S2). After examining the cross-correlation matrix among all metrics (presented in Supplementary Material S3), we selected MESH and NP landscape metrics for further analysis since they were among the least correlated and offered intuitive representations of connectivity and fragmentation dynamics (Jaeger, 2000).

We analysed the temporal dynamics of the NP and MESH metrics using time series functional principal component analysis (FPCA) implemented via the *fda* R package (Ramsay, 2024). Prior to FPCA, each time series was smoothed using B-splines with a basis size of five (nbasis = 5), employing the smooth.basis function. FPCA was then performed on the smoothed data to extract the dominant harmonics—i.e. independent temporal trends—underlying the time series. The first three principal components (harmonics) were retained to represent the main independent patterns of variation over time. Each time series received scores indicating how closely it resembled each of these principal harmonics. Supplementary Material S4 illustrates the shapes of the first three harmonics for both NP and MESH.

To further quantify trajectory characteristics, we explicitly derived the average slope of MESH and NP over time. For MESH, a linear model (lm) was fitted to the smoothed time series, while for NP, a second-degree polynomial model (poly,2) was fitted. This was because the former showed mostly linear declines, while NP changes were more non-linear and resembled a parabolic trajectory in many cases. In the latter case, slopes were calculated at each time step from the first derivative of the polynomial fit, and the mean slope across all time steps was used as the final indicator of average slope.

For all scenarios with a final habitat area of 20%, the NP trajectories were further clustered into three groups using time series k-means clustering with Euclidean distance as the similarity measure (Tavenard et al., 2020). This clustering allowed differentiation between simulations that were predominantly characterised by either habitat shrinkage with low fragmentation (minimal change in NP) or habitats exhibiting high fragmentation dynamics (high increase in NP).

### 2.4. Species metapopulation persistence

Species metapopulation dynamics were simulated using a novel individual-based model by Tao et al. (2024) that calculates metapopulation persistence in landscapes with different habitat configurations. Individual-based models of metapopulation dynamics provide a means to examine how species-specific traits (e.g. dispersal and competition) generate emergent patterns in species metapopulation persistence. This model was developed to identify how metapopulation dynamics augmented with competition can provide inferences on time until global extinctions, final abundances and spatial synchrony in species decline in response to landscape configuration (i.e. level of fragmentation). The model incorporated key species traits, namely mean fecundity, dispersal and density-dependent competition, to simulate the dynamics of individuals within landscapes composed of habitat and non-habitat classes. A parameter indicative of environmental stochasticity informing global reproduction success per generation was also used.

The individual-based simulation tracked all individuals within the two-dimensional binary habitat-matrix landscape over discrete time steps, where each time step indicates a generation of a species that reproduces once and then dies. The simulation began with an initial population of individuals randomly distributed across the landscape, retaining only those that fell within habitat locations. At each time step, individuals reproduce, and the offspring disperse across the landscape grid. The edges of the grid were connected top-to-bottom and left-to-right (wrapped around) to prevent dispersing individuals from leaving the landscape. Reproduction was governed by a Poisson process with a stochastic component, and dispersal followed a Gaussian distribution depending on the dispersal parameter *alpha*. Offspring that landed in non-habitat locations were removed, representing habitat filtering. The surviving offspring then underwent density-dependent competition, where local population density was convolved with a competition kernel to determine survival probabilities. Individuals in areas of lower competitive pressure were more likely to survive and form the next generation. The simulation proceeded iteratively until the entire population went extinct or a predefined time limit (500,000 time steps) was reached. Detailed mathematical formulations for these steps are provided in Supplementary Material S5.

In our simulations, most of the model parameters were adopted from Tao et al. (2024). Following Tao et al. (2024), the mean fecundity was set at 1.1, environmental stochasticity was set at 0.5 and competition weight (b) was set at 0.2. Both competition and dispersal processes were modelled using a Gaussian kernel with an exponent α of 1.5, which was the setting used to represent ‘resident’ species in Tao et al. (2024). We did not test additional dispersal parameters since an alpha of 1.5 is consistent with theoretical expectations of metapopulation dynamics, as proposed by Hanski (1998). Competition was modelled as a local density-dependent regulation, where individuals experienced competition-induced mortality based on the number of neighbouring individuals within the dispersal kernel (Tao et al., 2024). At the start of a simulation, individuals were placed randomly across a landscape with an initial population size of 5000 individuals, translating approximately to a mean density of 1.39 (5000/3600) individuals per cell. Individuals who were randomly placed in non-matrix pixels at this initial stage were immediately removed from the simulation. Notably, this population initialisation differed from that of Tao et al. (2024) since they used an initial population of 3600 instead of 5000, yet required the maintenance of species persistence in heavily fragmented landscapes over longer time steps.

Tao et al. (2024) ran metapopulation simulations on static landscapes. To integrate landscape change into the simulations, we incorporated a landscape update function that changes the landscape on which the simulation is run. This update occurred after each generation’s reproduction and dispersal phases, thereby ensuring that individuals had the opportunity for rescue dynamics immediately following landscape change (Fig. 1c). Landscape updates were triggered at three different rates relative to species generation time: once every generation (1:1), once every five generations (1:5), and once every 10 generations (1:10). We refer to this as the landscape update rate.

Each simulation proceeded until species extinction, which is defined as the loss of all individuals across the landscape. For each scenario, simulations were repeated 50 times to account for stochastic variability. The global extinction time (i.e. the number of generations from the start of the simulation until the population is extinct) was recorded, and the mean, minimum and maximum extinction times were computed for each landscape change and update rate scenario. To account for the skewed nature of the global extinction times, the log mean global extinction time is henceforth referred to as the species persistence.

### 2.4. Relating species persistence and landscape change

To assess potential relationships between species persistence and landscape change, we first produced box plots of species persistence across the different initial and final habitat percentage scenarios and landscape update rates. To analyse the impact of landscape change trajectories on species persistence, we ran random forest models on the FPCA scores of the first three principal components for both MESH and NP time series data.

We also modelled species persistence with only the final landscape values of MESH and NP using random forest models. These final habitat values were chosen because they represent current best practice in many landscape-ecological studies, where mostly only current-day landscape configurations are used to explain ecological processes. Comparing the fit of the models with the information from the trajectories of NP and MESH and those with the final landscape values allowed us to infer the importance of the trajectories of landscape change over and above the information derived from the static current conditions of the landscape. In addition to running the random forest models on all the decline scenarios together, we separately ran the random forest models on a habitat decline scenario with 20% final habitat (i.e.100%→20%, 75%→20%, 50%→20% and 25%→20%). Upon observing the final habitat percentage, these landscapes all look alike, differing only in their landscape change history. Thus, running the model on this subset allowed us to assess whether landscapes with the same final habitat area percentages show variation in species persistence that is correlated to the past landscape change trajectories (i.e. the history of the landscape).

To further investigate the shape of the landscape change trajectories and the impact on persistence, we used time series k-means clustering (Tavenard et al., 2020) to identify three different shapes of NP change and the related MESH changes over time. For the reasons provided above, we once again only used the landscape change scenarios with 20% final habitat in this analysis. To reduce the number of parameters affecting species persistence, we applied clustering to each initial habitat percentage group (i.e. 100, 75, 50 and 25%). For each such cluster, we then generated box plots of estimated species persistence at the three landscape update rates to infer the impact of varying landscape change trajectories.

Finally, we ran random forest models on species persistence using the ‘effective slope’ of NP and MESH (ΔNP or ΔMESH per generation) as explanatory variables. The effective slope was calculated by multiplying the average slope of each trajectory (from previous analyses) by the landscape update rate (1:1, 1:5 or 1:10). This formulation allowed us to express the rate of landscape change in terms of species generation time. For species with short generation times, many generations may occur before the landscape changes noticeably, whereas species with longer generation times may experience one or more landscape updates within a single generation. By expressing change per generation, the effective slope standardises landscape dynamics across species with differing life histories, allowing a consistent comparison of landscape impact on persistence.

We ran random forests on the species persistence with the effective slopes for each habitat decline scenario, starting from all four initial habitat percentages and ending at 20% of the total area. We note the accuracy (pseudo-R^2^), the variable importances and the partial dependence plots of the slopes from NP and MESH to draw inferences about which landscape metric is more influential in each scenario, along with understanding what the direction and range of influence is. The variable importance was calculated as the increase in the model’s Mean Squared Error (MSE) when the values of that variable were randomly permuted. For comparison, we also ran the models using the initial conditions of NP and MESH.

## 3. Results

### 3.1. Species persistence in different landscape change scenarios

Our results indicate that at lower landscape update rates (1:5 and 1:10), species tend to persist longer in landscapes with a larger historical habitat area. This can be observed by looking at the boxplots within the same final habitat percentage group, where, as the initial habitat percentage increases, the mean persistence time values also fall within higher ranges, with the highest median value for initial habitat percentage being 100%. This effect of past habitat—where the higher the past area, the higher the species persistence—becomes more prominent the slower the landscape changes (Fig. 1.c). This can be observed in the larger differences between median persistence for different initial habitat percentages in Fig. 1c, compared to Fig. 1a.

Overall, species persist longer in landscapes with higher final habitat percentages, as can be seen in Fig. 2.a–c, where the median values increase as the final habitat percentage increases to the right of the graph. However, within each group of landscape change scenarios resulting in the same final habitat percentage, the initial habitat condition continues to serve a significant role. In landscapes that change rapidly—at a rate comparable to species generation times (1:1 update rate)—the final habitat percentage has the most pronounced effect. In such cases, a larger final habitat area is strongly associated with greater species persistence, while the influence of past habitat areas becomes negligible.

**Figure 2:**
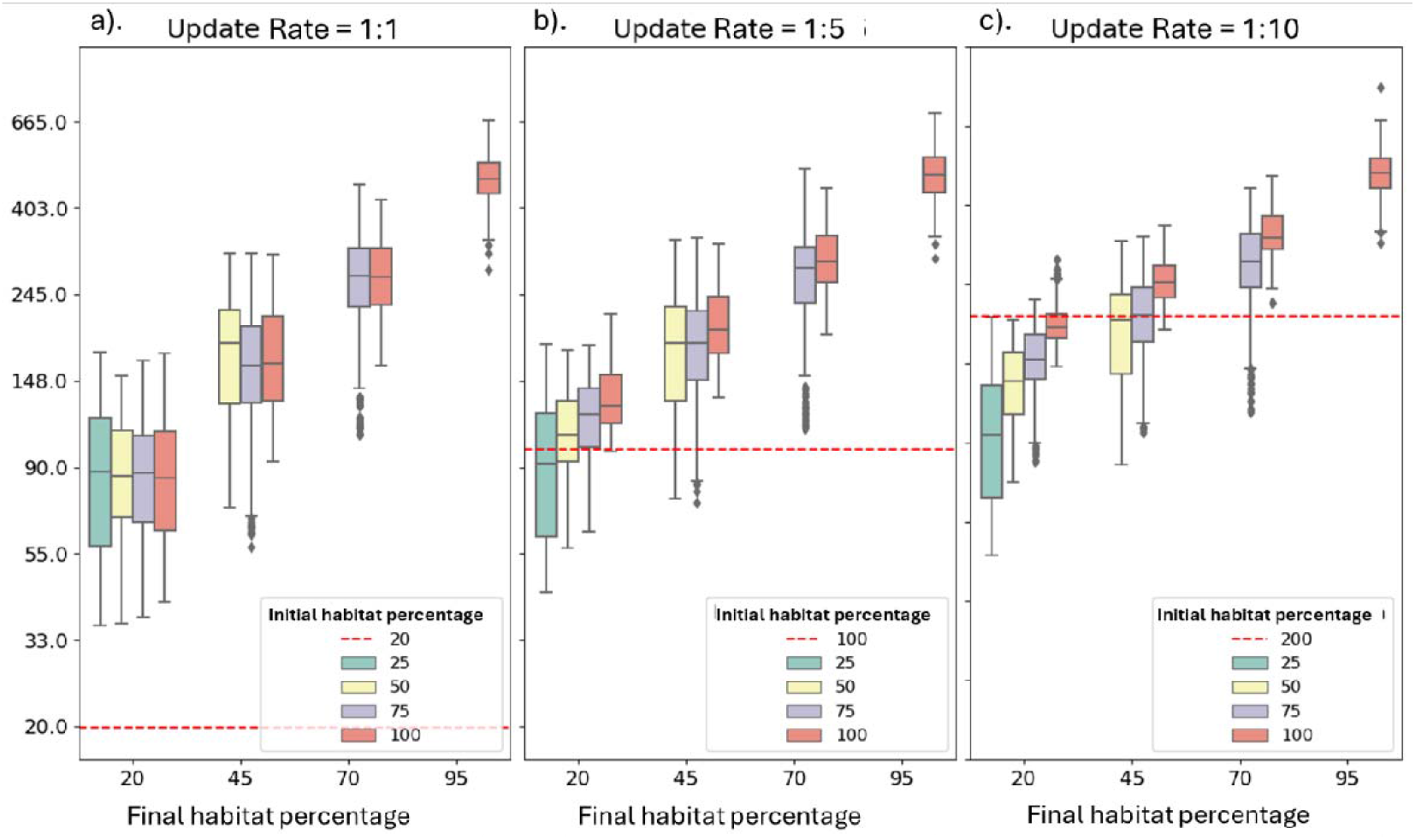
The distribution of mean species persistence times for different landscape change scenarios: a) rapid change, where the landscape updates with every species generation; b) medium change, where the landscape updates at every five species generations and; c) slow change, where the landscape updates every 10 species generations. Each graph is sorted from left to right as per the final habitat percentages of 20, 45, 70 and 95%. The colours of the boxplots indicate the initial habitat percentages. The red dotted line indicates the time step at which landscape change halts for each change scenario.

Notably, in slower-changing landscapes, the metapopulations often do not persist beyond the completion of the 20 time steps in the landscape change scenarios (red lines in Fig. 1). However, comparatively, the higher the initial habitat percentage in the simulated time series, the longer the persistence and the greater the likelihood of persisting beyond the end of the landscape change scenarios.

### 3.2. Effect of landscape change trajectories

At lower landscape update rates and for all scenarios that have a 20% final habitat percentage, the models with information on trajectories, which also inherently have information on both the initial and final conditions of the landscape, have a considerably higher fit (R^2^) than models with only final habitat condition information (Table 1). Specifically, when the landscape changes every 5 or 10 generations, the model with information on trajectories has R^2^ values of 0.64 and 0.77, respectively, whereas random forest models with only final landscape condition show much lower R2 values of 0.29 and 0.04, respectively. However, landscapes that change rapidly (i.e. with every generation of the species) show the best accuracy (56%) in the random forest with the final landscape conditions.

**Table 1:** Results of the random forest (RF) models fitted on scenarios with a final habitat percentage of 20%, as well as on all final habitat percentages. Columns show the speed of landscape change relative to the generation time of the species (landscapes updated every 1, 5 or 10 generations). Each block along the rows indicates variable sets used separately in RF models: a) RF model with time series trends, and b) RF model with initial and final time points. Green highlights show the two most important variables in each model.

| Landscape updates (every) | 1 generation<br>(Update rate= 1:1) |  | 5 generations<br>(Update rate= 1:5) |  | 10 generations<br>(Update rate= 1:10) |  |
| --- | --- | --- | --- | --- | --- | --- |
|  | 20% end<br>habitat | Full data | 20% end<br>habitat | Full data | 20% end<br>habitat | Full<br>data |
| *a) RF model with time series trends using FPCA* |  |  |  |  |  |  |
| R<sup>2</sup> | 0.51 | 0.87 | 0.64 | 0.88 | **0.77** | 0.88 |
| MESH harmonic PC1 | 2.48 | 24.67 | 6.47 | 35.13 | 14.41 | 28.11 |
| MESH PC2 | 2.24 | 24.09 | 2.28 | 16.11 | 2.71 | 7.88 |
| NP harmonic PC1 | 8.62 | 8.34 | 7.86 | 7.3 | 6.68 | 6.44 |
| NP harmonic PC2 | 3.37 | 2.91 | 2.79 | 2.63 | 3.07 | 2.84 |
| *b) RF model with only final habitat condition* |  |  |  |  |  |  |
| R<sup>2</sup> | **0.56** | **0.88** | 0.29 | 0.81 | 0.04 | 0.63 |

Among the time series variables, the PC1 components of both MESH and NP (see Supplementary Material S4) are the most important in determining species persistence. PC2 components primarily become relevant in models trained on the full dataset.

Compared to the random forest models that run on the scenarios with final habitat at 20%, the models that run on full data (all final habitat percentages) show higher R^2^ values across the landscape update rates. Here, MESH remained consistently more important than NP for the full data models, irrespective of the landscape update rate.

The time series clustering outcomes showed several trajectories for NP and MESH change over time given the same final conditions (20% habitat) and differing initial conditions. Even within the same initial conditions’ different trajectories of NP and MESH can be identified, due to the stochastic nature of the simulations. We determined that multiple distinct temporal trajectories of NP and MESH can emerge, even under scenarios with identical initial and final habitat amounts. These divergent fragmentation pathways, captured by time series clustering, reflect different structural responses to the same overall habitat loss. For example, in scenarios starting at 25% initial habitat (Fig. 3a), NP remains relatively stable across clusters, differing mainly in magnitude, while MESH declines in all cases but from different starting points. In contrast, when starting at 100% initial habitat (Fig. 3b), trajectories vary more in slope: one cluster shows a sharp rise in NP and steep decline in MESH (suggesting fragmentation), while another shows less increase in NP (indicative of shrinkage).

**Figure 3:**
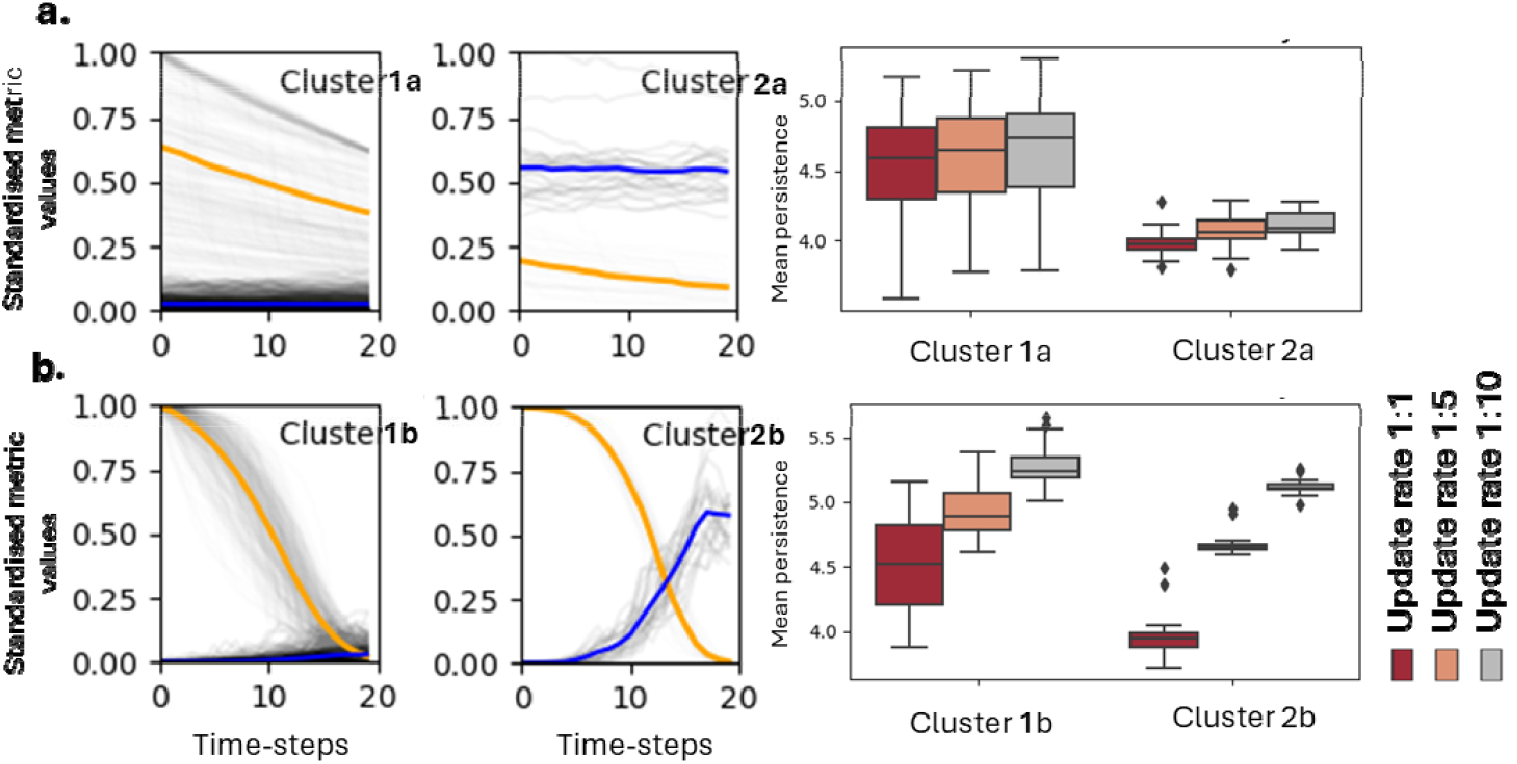
Left: trajectories of change over time for NP (blue) and MESH (orange; standardised per scenario between 0 and 1) for two time series clusters for landscape change scenarios with final habitat percentages of 20% and initial habitat percentage of (a) 25% and (b) 100%. Right: box plots pertaining to each cluster in each initial and final habitat percentage scenario; box plot colours indicate landscape update rates (see legend on the right).

We observed a much lower persistence in cluster 2a when compared to 1a, whereas this difference is less obvious between 1b and 2b, specifically for update rates of 1:5 and 1:10. Thus, systems that have fragmented from historically high habitat areas show lower variation in mean persistence if the landscape update rate is slow. The impact of the update rates of the landscape is also more evident in the 1b and 2b, with longer update rates showing higher mean persistence values. See Supplementary Material S6 for the graphs for all four initial habitat scenarios.

This differences in persistence due to landscape history can be significant. At the 1:10 update rate, the difference in the median persistence of scenarios with 100% initial habitat compared to those with 25% initial habitat can be more than 100 generations (cluster 2a versus cluster 2b; Fig. 3). In highly fragmented landscapes that started with 100% habitat (Fig. 3b, cluster 2b), the highest median persistence is approximately 180 generations (5.2 log mean persistence). In contrast, landscapes that began with only 25% habitat showed a low median persistence of approximately 65 generations (Fig. 3b, cluster 2a). At higher update rates, the influence of initial conditions on persistence time is lower. The differences in median species persistence between clusters 1a and 1b do not show such high differences (Fig. 3); however, the slower update rate shows considerably higher species persistence when the initial habitat percentage is high (Fig. 3b).

### 3.3. Influence of initial conditions versus the effective rate of change of the landscape

We determined that the predictors of species persistence vary depending on the initial habitat percentage. When the initial habitat is low (e.g. 25%), species persistence is best explained by the initial number of patches, with random forest models achieving an R^2^ of 0.75 (Fig. 4b). In contrast, when the initial habitat is high (e.g. 100%), the rate of change in habitat configuration—particularly the effective slope of MESH—becomes the strongest predictor, yielding an R^2^ of 0.82 (Fig. 4a). We observed a clear positive relationship between species persistence and both the effective slope and initial value of MESH, indicating that landscapes with higher MESH values and slower rates of decline tend to support longer persistence times (see partial dependence plots Fig. 4(a1), Fig. 4(a2). In the landscapes with 100% initial habitat, the absolute initial conditions (e.g. initial habitat area or patch count) have little explanatory power, while persistence is closely tied to how rapidly MESH and NP decline.

**Figure 4:**
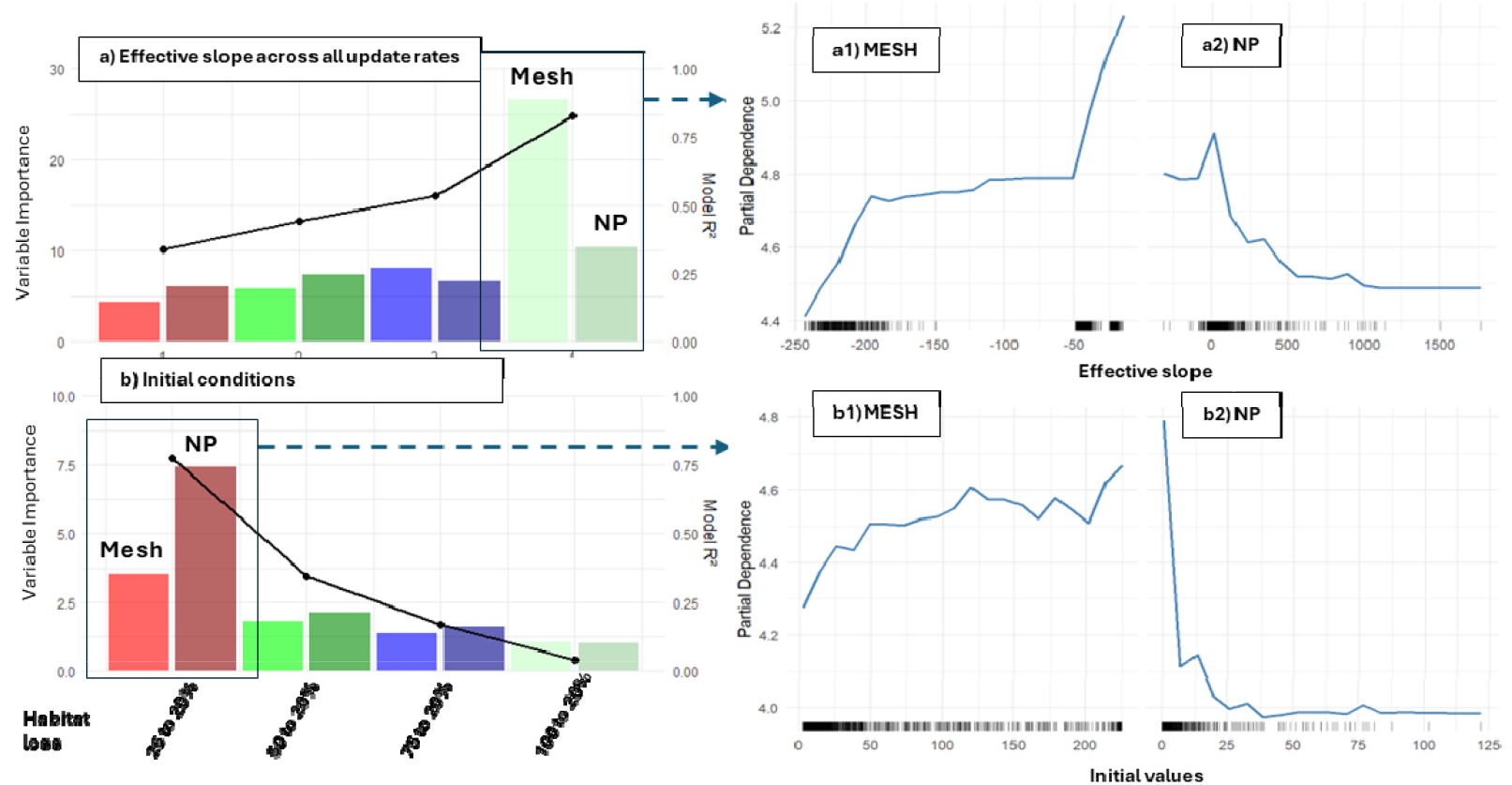
Top row: model results with the effective slope of change in effective mesh size (MESH) and number of patches (NP) (a). Bottom row: model results with initial values of MESH and NP. The first column indicates bar plots of variable importance and line plots of R^2^ values of the random forest. Each colour represents a habitat decline scenario (all final habitat percentages are at 20%) with differing initial conditions for habitat area (red: 25%; green: 50%; blue: 75%; sea green: 100%). The gradients of each colour indicate MESH (MS) and NP variables. Figures on the right show partial dependence plots for effective slope (a1,a2) and initial values (b1,b2) with respect to species persistence.

In contrast, in scenarios with minimal landscape change (e.g. a decline from 25 to 20%), NP metrics— particularly the initial number of patches—are more influential than MESH in explaining species persistence. This is partly because in low-change scenarios, initial NP values are often closely aligned with final habitat conditions, making them more predictive. Partial dependence plots (Fig. 4(b1), Fig. 4(b2)) show a clear, monotonically decreasing relationship between both the effective slope and initial NP values and species persistence. This suggests that landscapes with fewer, more aggregated patches tend to support longer species persistence than those with many small, fragmented patches.

## 3. Discussion

Our study provides a theoretical exploration of how long-term trajectories of landscape change influence species persistence. Three main findings emerge from our simulations. First, while the final habitat area governs the overall magnitude of persistence, historical habitat area significantly modulates it, particularly in gradually changing landscapes. Second, the speed of landscape change, scaled to species generation time, determines the strength and duration of time-lagged species responses, shaping whether species respond to current conditions or have a delayed response due to historical conditions. Third, the temporal pattern of landscape change, especially configurational changes, further influences the magnitude of persistence and time lags to extinction.

In support of the first finding, in landscapes with identical final habitat conditions, we observe that when landscape changes occur slowly (i.e. updates every 5 or 10 generations), species persist significantly longer when the initial habitat area is high. This reflects the influence of landscape history and aligns with the theory of extinction debt, where species continue persisting in deteriorated habitats due to delayed demographic collapse (Chen et al., 2023; Hanski and Ovaskainen, 2002; Kuussaari et al., 2009). In contrast, under rapid landscape change (updates every generation), persistence is explained almost entirely by final habitat area, indicating that when species turnover and landscape change occur at similar rates, species mainly respond to the current conditions (Watts et al., 2020). Roy et al. (2005) found that positive temporal autocorrelation can extend the persistence of metapopulations in degrading habitats, even in the absence of true source habitats, due to population inertia and the additional time it provides for potential recovery, temporarily creating source-like patches during decline. Therefore, since gradually declining landscapes tend to exhibit stronger temporal autocorrelation than those undergoing rapid decline, the mechanisms proposed by Roy et al. (2005) may help explain the delayed extinctions observed in our results. Notably, like Roy et al. (2005) we also only simulate sinking populations in patches, however, hypothesise that similar patterns can exist in source-sink dynamics also that are density capped.

A central contribution of this study is the demonstration that the rate of landscape change, scaled to species generation time (the effective slope), determines time lags (i.e. the delay between habitat loss and its impact on species persistence). In scenarios of severe habitat decline (e.g. 100% -> 20%), relatively slow declines in MESH and relatively slow increases in NP resulted in significantly longer time lags. These time lags, often exceeding 30 generations in our simulations, represent extended non-equilibrium states where species persist despite habitat degradation. While this study does not disentangle the precise mechanisms leading to these delays, it builds on other studies to assist in understanding how demographic and dispersal processes mediate delayed extinctions (Hylander and Ehrlén, 2013a; Loreau et al., 2003). In absolute terms, we estimate time lags of up to ∼1.5 log units (∼31 generations), which could imply decades of delayed extinction in unsuitable landscapes for species with annual turnover (Vellend et al., 2006). These findings underscore that species presence does not ensure long-term viability and that extinction risks may be significantly underestimated when based solely on current landscape metrics (Ridding et al., 2021).

While our model does not explicitly simulate species with varying generation times, we identify that species persistence is strongly correlated with the rate of landscape change per generation (i.e. how much environmental change a species experiences within its lifetime). For example, in a landscape that changes every five generations, both a long-lived species (e.g. 50-year generation time) and a short-lived species (e.g. 1-year generation time) would experience the same time lag in generational terms. However, under more rapid landscape change, long-lived species may endure multiple habitat shifts within a single generation, while short-lived species can respond across multiple generations through population turnover. In this sense, short-lived species may have more opportunities to compensate for change through dispersal and reproduction. To clarify, our model does not include internal demographic delays (e.g. multi-year reproductive windows or delayed maturity), which are important for understanding extinction lags in long-lived species (Jiménez-Franco et al., 2022; Watts et al., 2020). As such, our results are most applicable to species with short generation times and simple life histories (e.g. one reproductive cycle per generation), for which the timing of landscape change is more tightly coupled to ecological response. We hypothesise that in slowly changing landscapes, having more generations between landscape changes increases the likelihood of successful reproduction and dispersal. For species with limited dispersal, this may provide sufficient opportunity to persist in remnant habitat patches, even as overall fragmentation increases. Thus, species persistence may be more strongly influenced by the number of generations that occur between landscape changes than by the absolute length of each generation.

Beyond these core findings, we identify nuanced patterns in how landscape configuration affects persistence. The relative importance of connectivity (i.e. MESH) and fragmentation (i.e. NP) varies with the extent of habitat decline and initial habitat quantity. In scenarios with high initial habitat, MESH-based effective slopes best explain persistence, reflecting the role of large-scale connectivity loss (Graham et al., 2017; Jaeger, 2000). In contrast, NP becomes more informative in already degraded landscapes with low initial habitat, where limited change implies that the final configuration serves a greater role. This differentiation was supported by FPCA results, which showed that linear declines in MESH were consistent predictors of persistence under gradual change, whereas sudden increases in NP were more important in rapidly shifting landscapes (Semper-Pascual et al., 2021b).

We also determined that the shape of the trajectories of landscape change was either dominated by shrinkage (reduction from the edges, cluster 1b in Fig. 3) or fragmentation (splitting, cluster 2b in Fig. 3). We would like to reinforce that this definition of fragmentation is, as mentioned in Tao et al., (2024), technically fragmentation *per gradus*, which includes both area and connectivity loss in its definition. In our study, shrinkage and fragmentation had different effects on species persistence. In the 100% initial habitat scenario, shrinking landscapes (cluster 1b) marginally supported longer persistence compared to fragmenting ones (cluster 2b) when the update rate was low (1:5 or 1:10 generations). The low drop in persistence between shrinking and fragmenting clusters indicates that past spatial continuity and overall high habitat area can buffer against impacts of fragmentation. This is supported by the fact that fragmented landscapes with low initial habitat led to a much sharper drop in persistence when compared to less fragmented landscapes of similar habitat decline. These results suggest that fragmentation exerts time-delayed effects: species in once-continuous and large habitats may persist longer despite increasing fragmentation than those in landscapes that already historically had decreased habitat quantity (Krauss et al., 2010).

This study employed simulations of both landscape change and metapopulation dynamics to systematically investigate the influence of long-term trajectories of landscape change on species persistence. This approach offers a highly flexible parameter space by allowing the exploration of how different rates and magnitudes of long-term landscape change interact with metapopulation persistence. While our framework offers a theory-driven approach to understanding persistence dynamics, it simplifies certain ecological complexities. Assumptions such as uniform dispersal, wrap-around boundaries and fixed trait values may not fully reflect species-specific or real-world dispersal barriers (Justeau-Allaire et al., 2022; Tao et al., 2024). Moreover, the use of α = 1.5 to parameterise the dispersal kernel reflects species that have relatively short-distance dispersal and does not model species that migrate long distances. However, this setting proved to be useful for simulating metapopulation dynamics (i.e. asynchronous colonisation–extinction events that allow populations to persist despite local extinctions). Tao et al. (2024) concluded that ‘resident’ species (i.e. those with low dispersal abilities, as in our simulations) are more resistant to fragmentation than migrants (high dispersal). Based on the results of our study, it is evident that both dispersal abilities and landscape history influence resistance to fragmentation. In future research, interactions between landscape history and dispersal abilities could be analysed.

Due to our reliance on random fields and the use of linear decreases in habitat area to simulate landscape time series, non-linear change trajectories were not explored. The simulation of landscape change was mainly derived from spatially structured random fields, and habitat decline occurs uniformly, retaining the underlying spatial structure over time (Ovaskainen et al., 2002). It would be interesting to investigate how different spatial distributions of the stressors (e.g. roads or random perforations due to deforestation) further affect species’ persistence (Kun et al., 2019). However, our model is highly adaptable and can be extended to incorporate more complex trait distributions, empirical land use datasets or non-linear landscape transitions. Future work could also examine landscape change modes such as perforation or edge expansion (Miller-Rushing et al., 2019; Williams et al., 2021), and investigate how multi-species dynamics, trophic interactions and/or metacommunity processes shape persistence (Leibold and Chase, 2018; Zarnetske et al., 2017).

Overall, our results emphasise that delayed extinctions emerge from dynamic, trajectory-dependent processes governed by landscape history, configurational change and species life-history. By integrating species generation times with the temporal structure of landscape change, our framework helps identify when and for how long persistence may be delayed. From a conservation perspective, these delays represent both a critical opportunity to intervene before collapse and a risk, as apparent persistence can mask long-term vulnerability.

## Conclusion

Our study demonstrates that species persistence is not solely determined by current habitat area but is influenced by the rate, configuration and history of landscape change. Gradual habitat loss can create long extinction delays for landscapes in the same final habitat percentage, especially when the initial habitat area is high, while rapid change forces species to respond to present conditions. Landscape configuration, particularly effective mesh size and number of patches, further shapes persistence outcomes, with historical structure providing temporary buffers against decline. By linking landscape trajectories to species generation times, we highlight that delayed extinctions are a function of the effective rate of change in species-landscape systems. These findings emphasise the need to integrate the history of habitats into conservation planning since present-day species persistence may mask long-term vulnerability.

## Acknowledgements

We would like to acknowledge Alessandro Valentini for his effort in the initial methodology and software development. This research was funded by the Swiss National Science Foundation as part of the EMPHASES Project (Grant number: 200021_192018).

## Data Availability

The generated landscape data and simulation code has been uploaded to Zenodo: xxxxxxxxx.

## Declaration of generative AI and AI-assisted technologies in the writing process

During the preparation of this work the author(s) used ChatGPT 4.0 in order to correct the grammar and academic clarity of the writing. After using this tool/service, the author(s) reviewed and edited the content as needed and take(s) full responsibility for the content of the publication.

## References

Bertassello, L. E., Bertuzzo, E., Botter, G., Jawitz, J. W., Aubeneau, A. F., Hoverman, J. T., Rinaldo, A., & Rao, P. S. C. (2021). Dynamic spatio-temporal patterns of metapopulation occupancy in patchy habitats. Royal Society Open Science, 8(1). 10.1098/rsos.201309

Chen, X., Wang, Q., Cui, B., Chen, G., Xie, T., & Yang, W. (2023). Ecological time lags in biodiversity response to landscape changes. Journal of Environmental Management, 346(June), 118965. 10.1016/j.jenvman.2023.118965

Chen, Y., & Peng, S. (2017). Evidence and mapping of extinction debts for global forest-dwelling reptiles, amphibians and mammals. Scientific Reports, 7(March), 1–10. 10.1038/srep44305

Fletcher, R. J., Smith, T. A. H., Troy, S., Kortessis, N., Turner, E. C., Bruna, E. M., & Holt, R. D. (2024). The Prominent Role of the Matrix in Ecology, Evolution, and Conservation. Annual Review of Ecology, Evolution, and Systematics. 10.1146/annurev-ecolsys-102722-025653

Graham, L. J., Haines-Young, R. H., & Field, R. (2017). Metapopulation modelling of long-term urban habitat-loss scenarios. Landscape Ecology, 32(5), 989–1003. 10.1007/s10980-017-0504-0

Guardiola, M., Pino, J., & Rodà, F. (2013). Patch history and spatial scale modulate local plant extinction and extinction debt in habitat patches. Diversity and Distributions, 19(7), 825–833. 10.1111/ddi.12045

Hanski, I. (1998). Metapopulation dynamics. Nature, 396(6706), 41–49. 10.1038/23876

Hanski, I., & Ovaskainen, O. (2000). The metapopulation capacity of a fragmented landscape. Nature, 404(6779), 755–758.

Hanski, I., & Ovaskainen, O. (2002). Extinction debt at extinction threshold. Conservation Biology, 16(3), 666–673. 10.1046/j.1523-1739.2002.00342.x

Harisena, N. V., Grêt-Regamey, A., & Van Strien, M. J. (2024). Identification of metacommunities in bioregions with historical habitat networks. Ecology and Evolution, 14(8), 1–13. 10.1002/ece3.70076

Helm, A., Hanski, I., & Pärtel, M. (2006). Slow response of plant species richness to habitat loss and fragmentation. Ecology Letters, 9(1), 72–77. 10.1111/j.1461-0248.2005.00841.x

Herrault, P., Larrieu, L., Cordier, S., Gimmi, U., Lachat, T., Ouin, A., Sarthou, J., & Sheeren, D. (2016). Combined effects of area, connectivity, history and structural heterogeneity of woodlands on the species richness of hoverflies (Diptera□: Syrphidae). Landscape Ecology, 31(4), 877–893. 10.1007/s10980-015-0304-3

Hylander, K., & Ehrlén, J. (2013). The mechanisms causing extinction debts. Trends in Ecology and Evolution, 28(6), 341–346. 10.1016/j.tree.2013.01.010

Jaeger, J. A. G. (2000). Landscape division, splitting index, and effective mesh size: New measures of landscape fragmentation. Landscape Ecology, 15(2), 115–130. 10.1023/A:1008129329289

Jiménez-Franco, M. V., Graciá, E., Rodríguez-Caro, R. C., Anadón, J. D., Wiegand, T., Botella, F., & Giménez, A. (2022). Problems seeded in the past: lagged effects of historical land-use changes can cause an extinction debt in long-lived species due to movement limitation. Landscape Ecology, 37(5), 1331–1346. 10.1007/s10980-021-01388-3

Johst, K., Brandl, R., & Eber, S. (2002). Metapopulation persistence in dynamic landscapes: The role of dispersal distance. Oikos, 98(2), 263–270. 10.1034/j.1600-0706.2002.980208.x

Justeau-Allaire, D., Blanchard, G., Ibanez, T., Lorca, X., Vieilledent, G., & Birnbaum, P. (2022). Fragmented landscape generator (flsgen): A neutral landscape generator with control of landscape structure and fragmentation indices. Methods in Ecology and Evolution, 13(7), 1412– 1420. 10.1111/2041-210X.13859

Krauss, J., Bommarco, R., Guardiola, M., Heikkinen, R. K., Helm, A., Kuussaari, M., Lindborg, R., Öckinger, E., Pärtel, M., Pino, J., Pöyry, J., Raatikainen, K. M., Sang, A., Stefanescu, C., Teder, T., Zobel, M., & Steffan-Dewenter, I. (2010). Habitat fragmentation causes immediate and time-delayed biodiversity loss at different trophic levels. Ecology Letters, 13(5), 597–605. 10.1111/j.1461-0248.2010.01457.x

Kuipers, K. J. J., Hilbers, J. P., Garcia-Ulloa, J., Graae, B. J., May, R., Verones, F., Huijbregts, M. A. J., & Schipper, A. M. (2021). Habitat fragmentation amplifies threats from habitat loss to mammal diversity across the world’s terrestrial ecoregions. One Earth, 4(10), 1505–1513. 10.1016/j.oneear.2021.09.005

Kun, Á., Oborny, B., & Dieckmann, U. (2019). Five main phases of landscape degradation revealed by a dynamic mesoscale model analysing the splitting, shrinking, and disappearing of habitat patches. Scientific Reports, 9(1), 11149. 10.1038/s41598-019-47497-7

Kuussaari, M., Bommarco, R., Heikkinen, R. K., Helm, A., Krauss, J., Lindborg, R., Öckinger, E., Pärtel, M., Pino, J., Rodà, F., Stefanescu, C., Teder, T., Zobel, M., & Steffan-Dewenter, I. (2009). Extinction debt: a challenge for biodiversity conservation. Trends in Ecology and Evolution, 24(10), 564–571. 10.1016/j.tree.2009.04.011

Leibold, M. A., & Chase, J. M. (2018). Metacommunity ecology. Princeton University Press.

Liao, Z., Peng, S., & Chen, Y. (2022). Half-millennium evidence suggests that extinction debts of global vertebrates started in the Second Industrial Revolution. Communications Biology, 5(1). 10.1038/s42003-022-04277-w

Loreau, M., Mouquet, N., & Gonzalez, A. (2003). Biodiversity as spatial insurance in heterogeneous landscapes. Proceedings of the National Academy of Sciences of the United States of America, 100(22), 12765–12770. 10.1073/pnas.2235465100

Miller-Rushing, A. J., Primack, R. B., Devictor, V., Corlett, R. T., Cumming, G. S., Loyola, R., Maas, B., & Pejchar, L. (2019). How does habitat fragmentation affect biodiversity? A controversial question at the core of conservation biology. Biological Conservation, 232, 271–273. 10.1016/j.biocon.2018.12.029

Müller, S., Schüler, L., Zech, A., & Heße, F. (2022). GSTools v1.3: A toolbox for geostatistical modelling in Python. Geoscientific Model Development, 15(7), 3161–3182. 10.5194/gmd-15-3161-2022

Ovaskainen, O., Sato, K., Bascompte, J., & Hanski, I. (2002). Metapopulation models for extinction threshold in spatially correlated landscapes. Journal of Theoretical Biology, 215(1), 95–108. 10.1006/jtbi.2001.2502

Pan, Y., Hersperger, A. M., Kienast, F., Liao, Z., Ge, G., & Nobis, M. P. (2022). Spatial and temporal scales of landscape structure affect the biodiversity-landscape relationship across ecologically distinct species groups. Landscape Ecology, 37(9), 2311–2325. 10.1007/s10980-022-01477-x

Ramsay, J. (2024). fda: Functional Data Analysis (R package version 6.1.8). https://cran.r-project.org/package=fda

Ridding, L. E., Newton, A. C., Keith, S. A., Walls, R. M., Diaz, A., Pywell, R. F., & Bullock, J. M. (2021). Inconsistent detection of extinction debts using different methods. Ecography, 44(1), 33– 43. 10.1111/ecog.05344

Roy, M., Holt, R. D., & Barfield, M. (2005). Temporal autocorrelation can enhance the persistence and abundance of metapopulations comprised of coupled sinks. American Naturalist, 166(2), 246–261. 10.1086/431286

Semper-Pascual, A., Burton, C., Baumann, M., Decarre, J., Gavier-Pizarro, G., Gómez-Valencia, B., Macchi, L., Mastrangelo, M. E., Pötzschner, F., Zelaya, P. V., & Kuemmerle, T. (2021). How do habitat amount and habitat fragmentation drive time-delayed responses of biodiversity to land-use change? Proceedings of the Royal Society B: Biological Sciences, 288(1942). 10.1098/rspb.2020.2466

Slager, C. T. J., & De Vries, B. (2013). Landscape generator: Method to generate landscape configurations for spatial plan-making. Computers, Environment and Urban Systems, 39, 1–11. 10.1016/j.compenvurbsys.2013.01.007

Tao, Y., Hastings, A., Lafferty, K. D., & Ovaskainen, O. (2024). Landscape fragmentation overturns classical metapopulation thinking. Proceedings of the National Academy of Sciences. 10.1073/pnas

Tavenard, R., Vandewiele, G., Divo, F., Androz, G., Holtz, C., Payne, M., & Woods, E. (2020). Tslearn, A Machine Learning Toolkit for Time Series Data. Journal of Machine Learning Research, 21, 1–6.

van Strien, M. J., Slager, C. T. J., de Vries, B., & Grêt-Regamey, A. (2016). An improved neutral landscape model for recreating real landscapes and generating landscape series for spatial ecological simulations. Ecology and Evolution, 6(11), 3808–3821. 10.1002/ece3.2145

Vellend, M., Verheyen, K., Jacquemyn, H., Kolb, A., Van Calster, H., Peterken, G., & Hermy, M. (2006). Extinction debt of forest plants persists for more than a century following habitat fragmentation. Ecology, 87(3), 542–548. 10.1890/05-1182

Watts, K., Whytock, R. C., Park, K. J., Fuentes-Montemayor, E., Macgregor, N. A., Duffield, S., & McGowan, P. J. K. (2020). Ecological time lags and the journey towards conservation success. Nature Ecology and Evolution, 4(3), 304–311. 10.1038/s41559-019-1087-8

Williams, J. W., Ordonez, A., & Svenning, J. C. (2021). A unifying framework for studying and managing climate-driven rates of ecological change. Nature Ecology and Evolution, 5(1), 17–26. 10.1038/s41559-020-01344-5

Zarnetske, P. L., Baiser, B., Strecker, A., Record, S., Belmaker, J., & Tuanmu, M.-N. (2017). The Interplay Between Landscape Structure and Biotic Interactions. Current Landscape Ecology Reports, 2(1), 12–29. 10.1007/s40823-017-0021-5

