## Supplementary material for "Shape and rate of landscape change trajectories influence species persistence"

### Appendix

1. Details of matern variograms for simulating landscape change

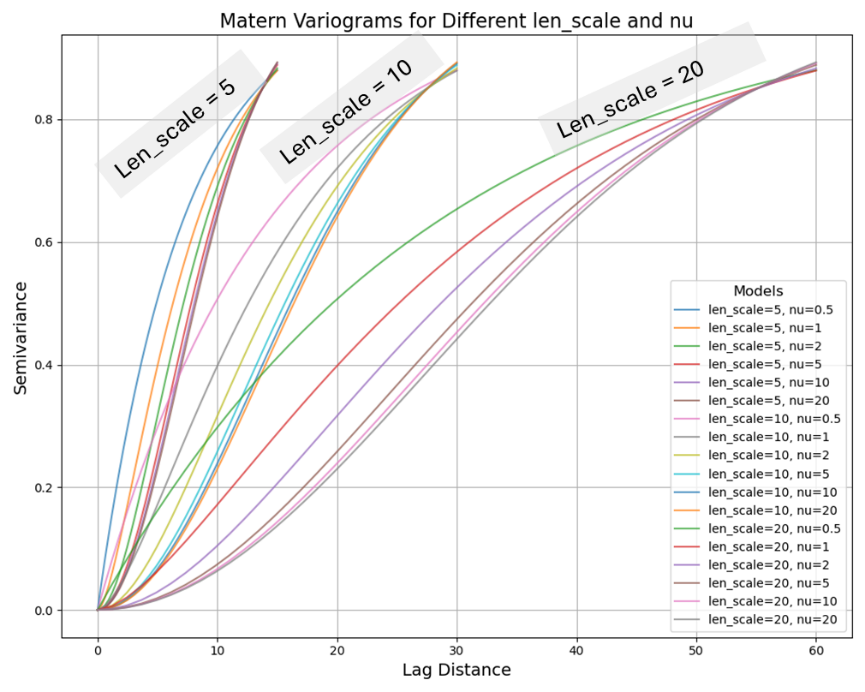

We simulate Matern-based Gaussian random fields based on:

$$Z(s)\sim GP(0,Matern(\nu=j\mathcal{,l=}i,\sigma2=1))$$

$where l\in\left\{ 5,10,20 \right\}$; $\nu\in\{0.5,1,2,5,10,20\}$

We transform the continuous-valued random field into a **binary habitat map** by thresholding:

For a given initial **habitat coverage** $A$%, and final habitat coverage of E%:

$$\tau_{I}={Percentile}_{100-A}(Z)$$

$$\tau_{E}={Percentile}_{100-E}(Z)$$

Then define :

$$H_{I}\left( s \right)= \left\{ \begin{aligned} 1, Z\left( s \right)\geq\tau_{I} or \tau_{E} \\ 0, Z\left( s \right)< \tau_{I} or \tau_{E} \end{aligned} \right.$$

This results in a **binary matrix** $H_{I}\left( s \right)\in\left\{ 0,1 \right\}n\times n$ representing habitat presence (1) or absence (0) and $n$ is the dimensions of the square lattice. In this case 60 by 60 pixels.

Time steps of landscape in between the initial and final habitat percentages is based on τ chosen as *x* equally spaced percentiles between the initial and final percentile values where *x* defines the number of time-steps, and the transition from τ_A_ to τ_E_ models habitat degradation or connectivity changes.

We estimate 20 such equally spaced thresholds, and compute 20 landscape steps between a set initial and final la

2. Simulation trajectories of landscape change scenarios between 4 landscape initial habitat area percentages (100,75,50,25%) and 4 remaining habitat percentages (95,70,45,20%), colours indicate scenarios with similar remaining habitat percentages.

Area change

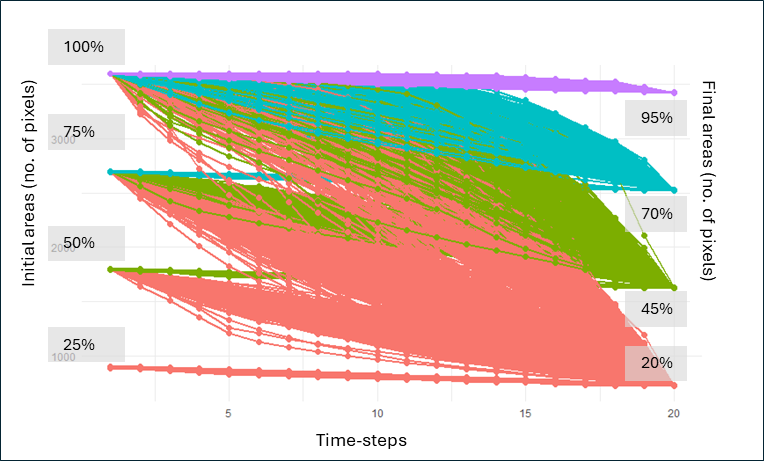

Effective mesh size change

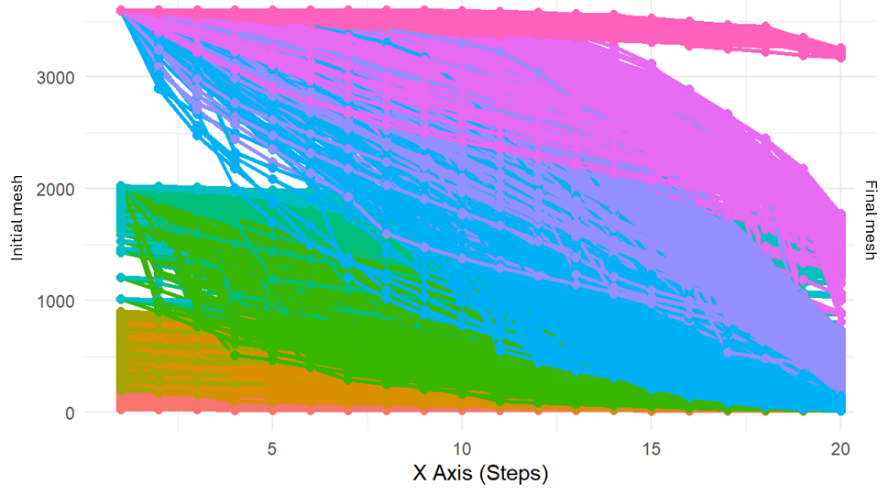

No. of patches change

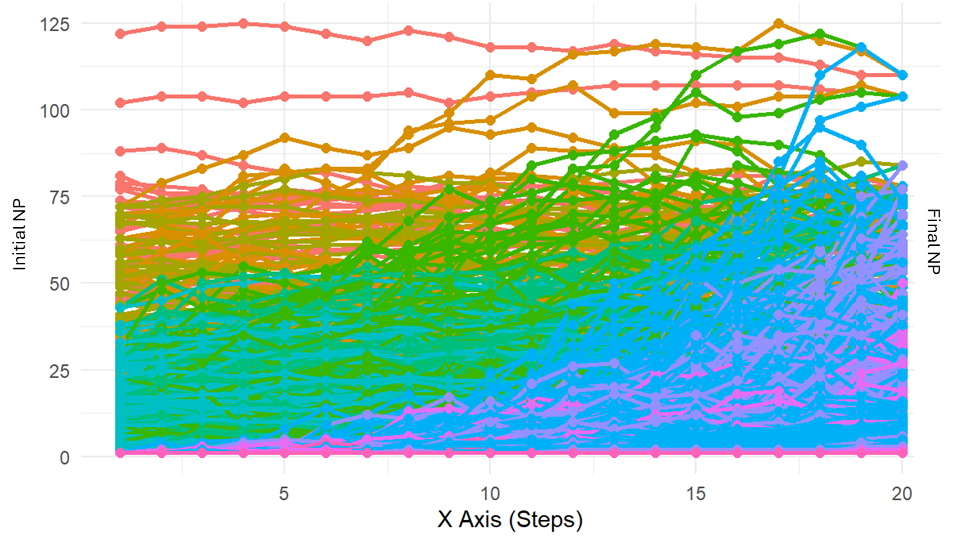

3. Correlation matrix for the computed time-series of landscape metrics. Data was grouped into two similar groups. A) with MESH and Area and all metrics correlating to these over time (slope) and initial values; and B) correlating with NP.

A)
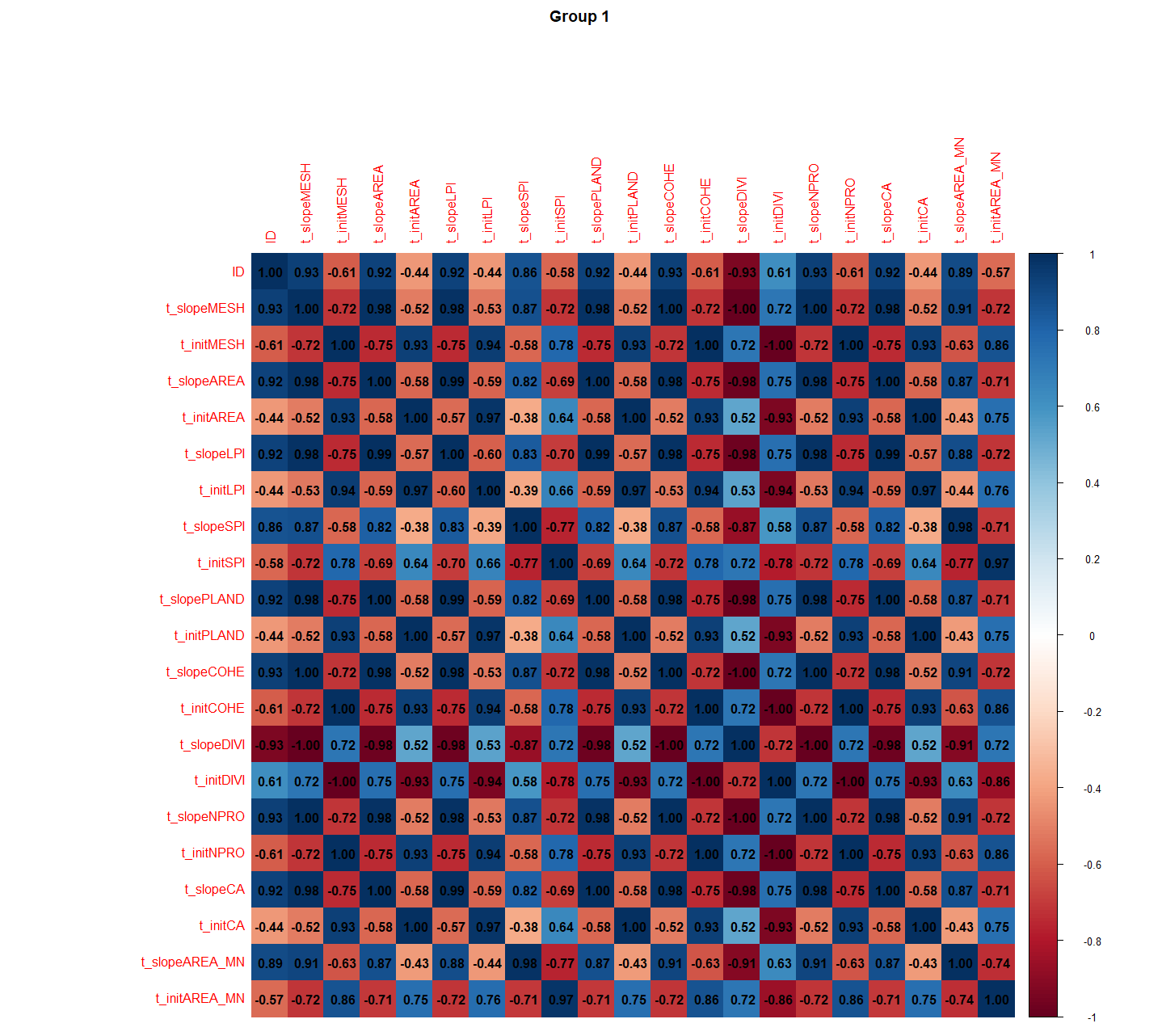

B)
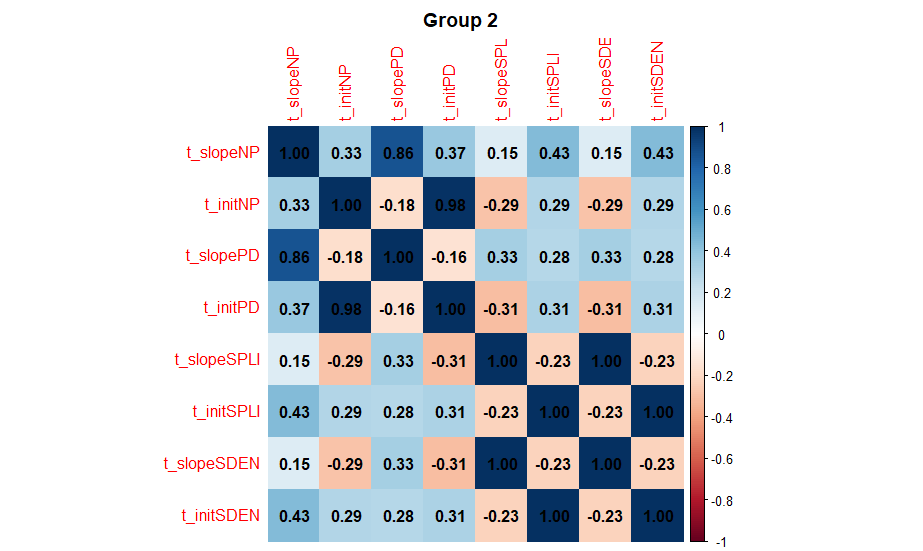

4. Functional Principal Component Analysis to identify principle harmonics in landscape metric change over time

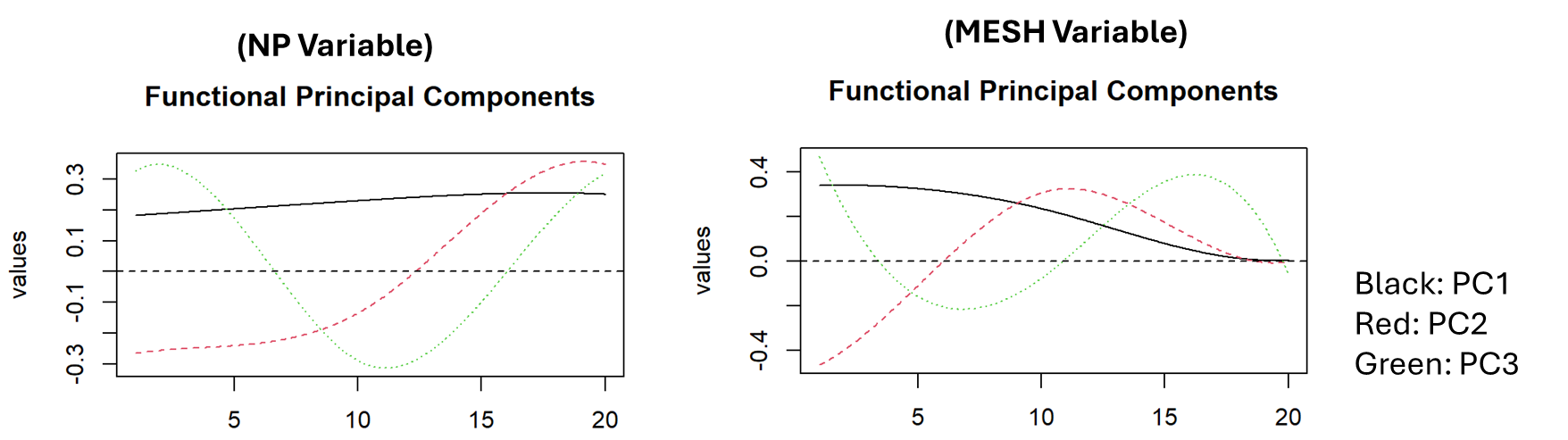

5. Individual Based Models are algorithmic models that in our case work on these steps (as developed by (Tao et al., 2024):
a. **Species dynamics:**

**Step 1:** Initial population locations are drawn from a uniform random distribution between (0,size of landscape (S)] :

$$\left( x,y \right)\sim{\mathrm{Uniform}\left( 0,1 \right)}^{2}S$$

**Step 2:** Cross check if the coordinates of the initial population coincide with binary habitat landscape. All randomly chosen coordinates overlapping with value 1 for habitat gets stored as a species location as is defined as the *parents*.

**Step 3**: Each parent $i$ reproduces a number of offspring ($k_{i}$) based on a draw from a Poisson distribution with parameter $\mu$ _RS_ that defines a global the reproduction success rate ( $\mu$ _RS_) for the generation. This resproduction success rate is defined by a fecundity rate ($\mu$) and an environmental stochasticity parameter ($\sigma_{R}$) based on:

$$\mu_{RS}= LogNormal(\log\left( \mu,{\sigma_{R}}^{2} \right))$$

And the actual number of offsprings$k_{i}$ per parent $i$ is estimated as:

$$k_{i}\sim Poisson(\mu_{RS})$$

**Step 4**: The offsprings disperse away from the parent locations to new coordinates $\left( x^{'},y^{'} \right)$based on random draws from a gaussian kernel for a set dispersal threshold ($\alpha$):

$$\Delta x,\Delta y\sim N\left( 0,\alpha^{2} \right)$$

$$\left( x^{'},y^{'} \right)=\left( x\boldsymbol{+}\Delta x,y\boldsymbol{+}\Delta y \right)$$

In our case we used $\alpha=1.5$, implying that the individual has a high probability of landing within 1.5 pixels of the original pixel, indicating small-scale dispersal.

**Step 5:** Cross check whether these new locations coincide with binary habitat landscape. All locations overlapping with value 1 (habitat) only gets stored (overwrites) as the set of offspring locations.

**Step 6:** Once the offspring reaches the destination cell $\left( x^{'},y^{'} \right)$ it faces a density dependant competition in that location. The density dependence is defined by a gaussian competition kernel (K) that scales based on the dispersal distance threshold ($\alpha)$ i.e.:

$$K(x,y)\propto\exp^{(-\frac{\left( x^{2}+y^{2} \right)}{2\alpha^{2}})}$$

This competition kernel is convolved with the density of offsprings (D) in each location in the landscape giving rise to a density weighted local competition that estimates that at each location (x,y) what is the total competitive pressure from all individuals in the surrounding area, weighted by distance:

$$C(x,y)=(K*D)(x,y)$$

The offsprings in each location then accrue a survival probability based on a mortality rate (m) as defined by this density dependent competition and a parameter ‘b’ that defines the strength of this competition bounded between 0,1:

$$m=1- \frac{1}{1+b\cdot C(x,y)}$$

Next again for each offspring a random number is drawn from a uniform distribution between 0 and 1. If the random draw is greater than the mortality rate then the offspring survives. All survived offspring locations are again stored (overwrites) as the set of offspring locations.

**Step 7**: The algorithm now attributes these survived offsprings locations as the new parent locations and steps 3-6 are repeated until no offspring remains. Each 3-5 step represents one generation time of the species.

For each dispersal step there is a mod function applied to the output location that allows the dispersal destination to be “wrapped around” the landscape i.e. returns into the landscape from the opposite side if destination falls outside the extent.

b. **Landscape dynamics:**

Into this we load the landscape update rule such that after each generation reproduces, they disperse into a new landscape that has declined from the previous one. This change occurs for 20 time steps and details of this is given in S1.

6. Temporal clusters of landscape configuration change for all four scenarios of initial habitat (row-wise: 100%,75%,50% 25%) and ending in 20%. Blue lines show profile in NP and orange lines show profile of Mesh. Right column shows persistence bar plots for each cluster and three update rates.

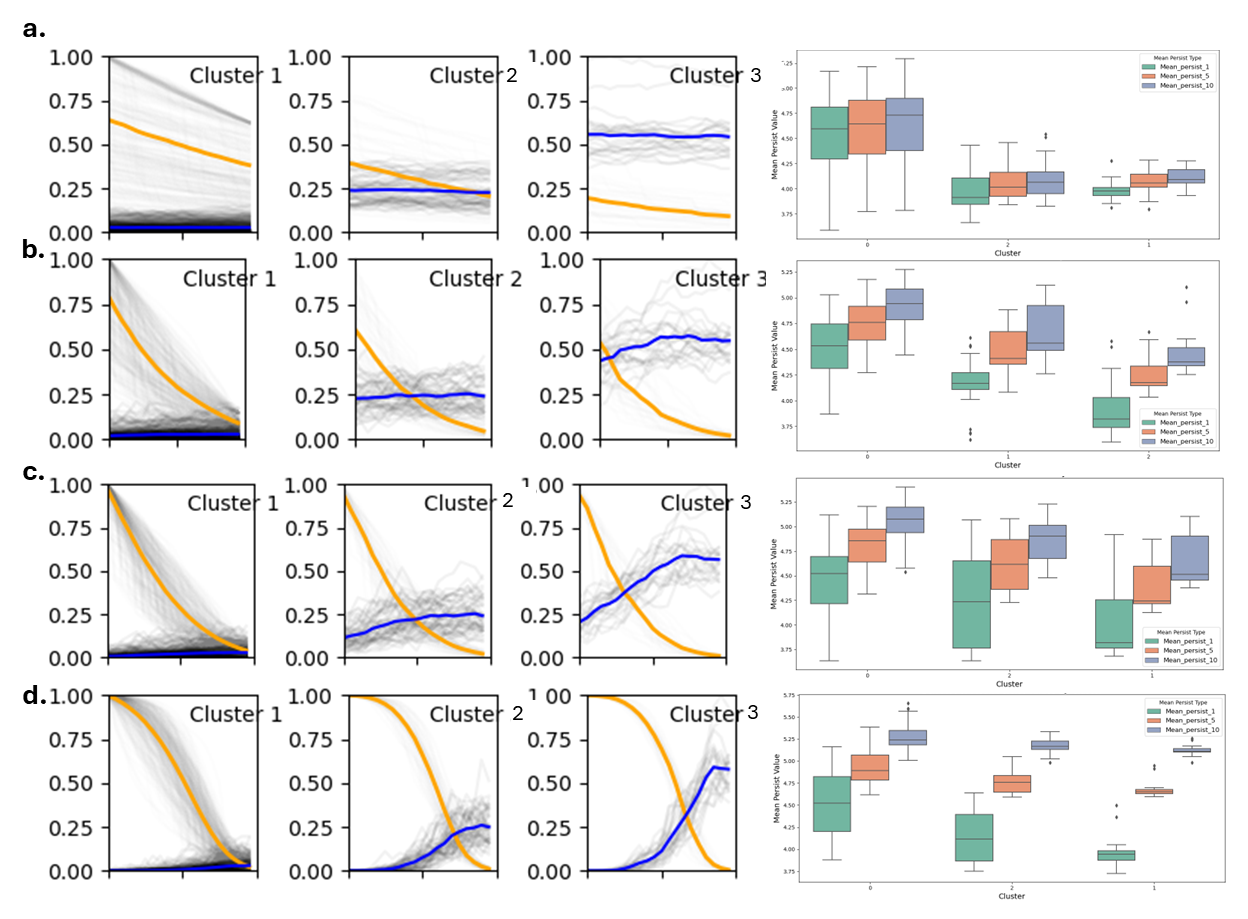
